# Inoculation with arbuscular mycorrhizal fungi can increase the concentration of PFAS compounds in cereal crops

**DOI:** 10.1101/2025.09.09.674819

**Authors:** Stephanie J Watts-Williams, Sara Thomas, Alison R Gill, Thi Diem Nguyen, Shervin Kabiri

## Abstract

Contamination of agricultural soils with per and poly-fluoroalkyl substances (PFAS) is now widespread and common throughout the world due to management practices such as biosolid application. Crop and pasture plants can readily take up and accumulate PFAS in their tissues, which eventually ends up in food products or livestock feed and presents health issues. We investigated whether arbuscular mycorrhizal (AM) fungi in the soil affect plant PFAS uptake in three important cereal crop species (barley, bread wheat, durum wheat), using a combination of soils that were artificially-spiked or naturally contaminated with PFAS. Spiking soil with PFAS did not interfere with AM colonisation of roots, nor did it detrimentally affect plant biomass. However, the shoots of all three plant species accumulated high concentrations of multiple short- and long-chain PFAS compounds. Inoculation with AM fungi led to increased concentration of all measured PFAS compounds in durum wheat shoots and several compounds in barley shoots, with no effect on any compounds in bread wheat shoots. The effect of AM fungal colonisation on PFAS concentrations in cereal plants is highly dependent on the nature of the soil PFAS contamination (i.e., the concentration and speciation) and the host plant species. It remains to be investigated whether AM fungi can directly take up PFAS from the soil.

## Introduction

Per and poly-fluoroalkyl substances (PFAS) are a group of highly stable chemicals that are common components of household and industrial products including adhesives, coatings, packaging, and pesticides since the 1940s (Wang et al., 2020). PFAS do not easily degrade in the environment and persist in terrestrial and aquatic systems alike. This allows PFAS to easily enter the food chain at several different points in our food production systems, bioaccumulating in plants, livestock and eventually humans. The accumulation of PFAS in human bodies can lead to serious health issues including immune and endocrine disorders, cardiovascular disease, and reproductive and developmental disruptions (Jha et al., 2021). The cost of PFAS contamination in the food chain to the health system is estimated at €52– 84 billion per annum in Europe alone (Cordner et al., 2021), so the mechanisms for PFAS uptake from soil and water into food crops must urgently be uncovered in order to devise ways to reduce or block bioaccumulation at this step in the food chain.

Agricultural soils become contaminated with PFAS through a number of pathways including contaminated irrigation water, application of organic matter to the soil as an alternative to conventional fertilisers (e.g., sewage sludge, biosolids), leaching from landfill, and pesticide use (Wang et al., 2020). Plants, including many popular food crops, readily take up the small and water-soluble PFAS via their roots, and accumulate them in shoots and grains (Adu et al., 2023). A study by Ofoegbu et al. (2022 in wheat plants grown in soil spiked with perfluorooctane sulfonic acid (PFOS) to different concentrations found that, although PFOS contamination did not affect the yield of the plants, PFOS accumulated substantially in the wheat grains, and led to a reduction in concentration of essential plant elements (phosphorus, potassium and magnesium). Meanwhile, Wen et al. (2014 conducted a field study that evaluated increasing applications of biosolids on the uptake and translocation of PFAS in wheat plants, and reported that the transfer factors of PFAS compounds from the straw to the grain was higher for perfluorosulfonic (PFSA) than perfluorocarboxylic (PFCA) acids.

A relatively unexplored avenue for the uptake of PFAS by plants is that of arbuscular mycorrhizal (AM) fungi. AM fungi are ubiquitous in agricultural soils (Frew et al., 2025), and benefit plant growth by acquiring inorganic nutrients and other solutes from the soil and transporting them into the plants via structures in the root cortical cells called arbuscules (Smith and Read, 2008). Several studies are showed that AM fungi can reduce the uptake of soil contaminants such as heavy metals into the host plant, conferring a positive effect on plant biomass compared to a non-colonised control plants (Chen et al., 2003; Watts-Williams et al., 2013). This is known as a ‘protective’ function of AM fungi that contrasts with the function of soil nutrient uptake that the association is typically reported in the literature. A recent experiment found that soil perfluorooctanoic acid (PFOA) contamination led to reduced AM colonisation in spring onion plants, and negatively affected plant performance (as biomass and root morphology) (Yan et al., 2025). The authors concluded that inoculation of the plants with AM fungi may have reduced the negative effect of PFOA contamination on the host plant, through alterations to resource allocations strategy that favours shoot growth. It has also been demonstrated that AM fungi interact with the plant uptake of microplastics, another emerging soil contaminant (Chen et al., 2023); while the total amount of poly methyl methacrylate (PMMA) taken up by the plant was not reduced, the translocation of PMMA from roots to shoots was significantly reduced by AM colonisation, including evidence that PMMA particles were physically immobilised inside AM fungal vesicles and intraradical hyphae.

The effect of AM fungi on PFAS uptake or translocation in plants has not yet been explored. Given there is no existing data, there are three possible hypotheses regarding a potential effect of AM fungi on plant tissue PFAS concentrations:

1. There is no effect of AM colonisation on plant PFAS concentrations;
2. AM fungi ‘protect’ the host plant and reduce the concentration of PFAS in shoots; or
3. AM fungi promote host plant uptake of PFAS from the soil and thus increase PFAS concentration in shoots.

Once we have an understanding of how AM fungi affect plant PFAS uptake, we can target future research to best manage AM fungi and soil biology more broadly, while minimising PFAS uptake into crops. This may be done through introducing new species of AM fungi into fields that best immobilise PFAS to prevent plant uptake, by selecting crop varieties that do not associate with AM fungi to suppress PFAS uptake, or by using AM fungi to facilitate plant-based strategies for soil remediation (phytoextraction) (Kavusi et al., 2023).

The following experiments had the overarching aim to determine the effect of soil PFAS contamination and AM fungi on three important cereal crops (barley, bread wheat, and durum wheat), with the specific aims to determine:

i. Whether there is interspecific variation in plant response to PFAS contamination;
ii. Whether soil PFAS contamination affects the potential for AM fungi to colonise roots; and
iii. Whether inoculation with AM fungi increases resilience of host plants to PFAS contamination in the soil.

## Methods

### Soil preparation and AM fungal inoculation

The soils used were a sandy soil collected from a PFAS-uncontaminated area in South Australia (Soil 1), and a sandy soil from an aqueous film-forming foam (AFFF)-contaminated area (Soil 2) due to the repetitive application of the AFFF over time. Soil 1 had no PFAS contamination; half of Soil 1 was spiked with AFFF to a ΣPFAS of 6.9 mg kg^-1^ and the other half was not spiked (control). Soil 2 had an existing ΣPFAS of 1.9 mg kg^-1^ soil, which comprised different types of PFAS (Table S1. Supporting Information) while PFOS was the dominant compound (1.76 mg kg^-1^). Soil 1 had pH value of 8.3 and Soil 2 had pH of 7.2. Plant-available (Colwell) phosphorus was <5 P kg^-1^ for both soils, and nitrate was 6.7 and 1.0 mg N kg^-1^, and ammonium <1 and 2.1 mg N kg^-1^. Soils were sieved to <2 mm and then Soil 2 was mixed with a fine beach sand to dilute the existing PFAS concentration, while Soil 1 was mixed approximately 50:50 *w/w* with the fine beach sand to match the texture of the Soil 2. The final concentration of PFAS in the spiked soil (Soil 1) was greater than the AFFF-contaminated soil (Soil 2), while unspiked soil (Soil 1) had no PFAS. Each 100 mm squat punnet pot held 500 g of soil/sand mix.

### Experiment 1 – supplementation of the native AM fungal community with a commercial inoculant

To test the effect of introduced AM fungi to field soils containing the native AM fungal community, both Soil 1 and Soil 2 were inoculated with a commercial product from MicrobeSmart (StartUp Ultra) that contained four isolates of the AM fungus *Rhizophagus irregularis*. Average AM fungal spore number was estimated by sieving and microscope, and the product was added at a rate of ∼400 spores plant^-1^ to the +AMF treatment soil. No spores were added to the non-inoculated (control) soil.

### Experiment 2 – inoculation of sterilised soil with AM fungi

To test the effect of AM fungal inoculation on plant PFAS accumulation against a truly non-mycorrhizal control (ie., no native AM fungi present), Soil 1 was autoclaved two times prior to inoculation with AM fungi (conducted as for Expt. 1).

### Plant preparation and growth conditions

In both experiments, three species of cereal crop were included: *Hordeum vulgare* (barley) var. Compass, *Triticum aestivum* (bread wheat) var. Scepter, and *Triticum durum* (durum wheat) var. Westcourt. The species and varieties were selected after consultation with the relevant industry, based on their popularity in Australian agriculture. For both experiments, all seeds were surface sterilised in 10% bleach solution for 10 minutes, rinsed well with RO water, then moved onto moist filter paper and sealed in petri dishes, before being left at room temperature under laboratory lighting to germinate, with occasional watering when the filter paper was almost dry. Three days later, the germinated seedlings were transplanted to the prepared soils. There were five biological replicates of each treatment.

All plants were grown in a controlled environment chamber where the day and night temperatures were 24 ºC and 18 ºC, respectively, and daylength was set at 16 hours. Plants were watered 3-4 times per week to 15% of the soil mass (i.e., 60 mL available water). Plants were given 10 mL of full-strength Long-Ashton nutrient solution on three occasions during the growing period.

### Harvest and sample analysis

Plants were destructively harvested 73 days after transplantation to the soils. A 10 g subsample of soil was preserved from each pot. Roots were washed free of soil using RO water, then shoots and roots were separated and weighed. A subsample of fresh roots of approximately 200 mg was taken from each plant and submerged in 50% ethanol solution for root staining and AM fungal colonisation quantification. After sub-sampling, the roots were weighed again and then the root and shoot samples were placed in 25 mL plastic sample jars and covered with a Kimwipe secured by rubber band, in preparation for freeze-drying. All fresh plant samples were then freeze dried for 72 hours at −20 ºC. Following drying, the plant samples were weighed again to obtain shoot and root dry weight data.

### Plant PFAS analysis

PFAS was extracted from the plant tissues following modified methods by Nassazzi et al. (2022). The plant samples were freeze dried and then 0.1 g of the dried samples homogenised in ball mill (Retch) using a zirconium jar and balls using three balls in 100 mL jar. Each milling conducted for 30 seconds at 500 rpm and after milling the jar and balls were washed with water first followed by LC-MS grade methanol to avoid any PFAS contamination to next samples. All samples were spiked with PFAS stable isotope solution at 5 ng mL^-1^ in the final extract. The sample was vortexed and equilibrated for 30 minutes prior to extraction, then extracted three times with 4 mL of methanol containing 0.1 M of ammonium hydroxide (NH_4_OH). Each extraction consisted of 5 minutes of vortex followed by centrifugation for 10 min at 2500 g. Supernatant from the three extractions were combined and evaporated under N_2_ in a 40 °C using a Dry Block Multivap Evaporator to 1 mL. The extracts were then passed through an SPE cartridge (Agilent, Bond Elut Carbon) which was pre-conditioned with methanol. The final eluent was collected into a polypropylene autosampler vial, blown down under N_2_ and made to a final volume of 1 mL. A spiked control sample and a method blank were extracted alongside each batch of samples.

Chromatography was performed on an Agilent 6495 liquid chromatography/triple quadrupole mass spectrometer (LC/TQ) system. The separation of PFAS compounds was done on an Agilent ZORBAX RRHD Eclipse Plus C18 (2.1 × 100 mm, 1.8 µm), with an Agilent guard column ZORBAX RRHD Eclipse C18 (2.1mm, 1.8 µm), equipped with an InfinityLab PFAS delay column, 4.6 × 30 mm. Mobile phases were 2 mM of ammonium acetate in 95% water, 5% acetonitrile (A) and 100% acetonitrile (B). Injection volume was 5 µL (Agilent 1290 Infinity II) and the flow rate was 0.4 mLmin^-1^. The column oven was kept at 40 °C and the autosampler at 10 °C. The solvent gradient and LC/TQ instrument parameters are provided in Table S2 and S3 of Supporting Information. Negative electrospray ionization was used with calibration range of 0.01 to 100 ng mL^-1^. All standards contained the same 13C PFAS concentration as the samples for each run. Every 10 to 15 samples, a solvent blank and a standard solution were analysed to track instrument performance. Calibration curves weighed 1/x, automated peak integration was used however, integrations were manually curated to ensure accuracy.

Agilent 6495 LC/TQ with Agilent Jet Stream (AJS) electrospray ion source was operated in dynamic multiple reaction monitoring (dMRM) mode. This allows the addition of more MRM transitions for additional compounds if needed. The LC/TQ autotune was performed and all data acquisition and processing were performed using the Agilent MassHunter software version 10.

### AM colonisation analysis

Root samples preserved in ethanol were rinsed thoroughly with RO water before being placed in a 10% KOH solution for seven days at room temperature. Cleared roots were rinsed well with RO water before being submerged in a 5% ink in vinegar solution at 60°C for 15 minutes (Vierheilig et al., 1998). The stained roots were subsequently rinsed thoroughly and allowed to de-stain in a 5% vinegar solution for one day, before being moved to a 50% glycerol solution for storage and microscope analysis. Root length colonised by arbuscular mycorrhizal fungi was quantified by the gridline intersect method (Giovannetti and Mosse, 1980).

### Statistical analysis

All data were checked for normality using the Shapiro-Wilk test. Any variables that did not conform to the assumption of normality (*P*>0.05) were analysed using the Kruskal-Wallis test instead of Student’s t-test.

Expt.1 plant data included AM colonisation, biomass and shoot PFAS concentrations (sulfonic acids), which were analysed by two-way ANOVA where the factors were *Mycorrhiza* and *Soil*, for each plant species, respectively. Only the PFAS-contaminated soils (Soil 1 spiked and Soil 2) were analysed for shoot PFAS concentrations (sulfonic acids), and a Student’s t-test was employed for each compound to compare concentrations in the AM-inoculated vs non-inoculated plants of each plant species, respectively.

Expt. 2 plant biomass data were analysed by two-way ANOVA with the factors *Mycorrhiza* and *Soil*; only the spiked soil plants were analysed for tissue PFAS concentrations (sulfonic and carboxylic acids), and Student’s t-test was employed for analysis, with *Mycorrhiza* as the factor.

A principal components analysis (PCA) was undertaken using the “PCA” function in the “FactoMineR” package, including all the shoot PFAS compounds concentrations from the Expt. 2 plants. The PCA biplots were drawn using the “factoextra” package and the scores coloured by Crop; the group mean was also computed for each level, and a 95 % confidence ellipse drawn around the mean to determine significant differences between groups.

All statistical analyses were conducted and graphs plotted using R Statistical Software.

## Results

### Experiment 1

#### AM colonisation

All the plants were well-colonised by AM fungi, with root length colonised values ranging from 40.4% (durum wheat, Soil 1 spiked) to 99.3% (barley, Soil 1 unspiked) (Figure 1). In this study, there was little evidence of a detrimental effect of PFAS on AM colonisation of roots – only in barley, and only a small reduction in colonisation percentage. For bread wheat, the plants grown in Soil 2 were more colonised by AM fungi than in Soil 1 (see ANOVA outcomes in Table S4 Supporting information). In durum wheat, a similar result was observed, where the plants grown in Soil 2 were generally more colonised by AM fungi than in either treatment with Soil 1. In barley, root AM colonisation was effectively saturated across all treatments (mean value for barley was 96.6% root length colonised), but AM colonisation was significantly higher in the unspiked Soil 1 plants compared to the spiked Soil 1 plants.

**Figure 1.**
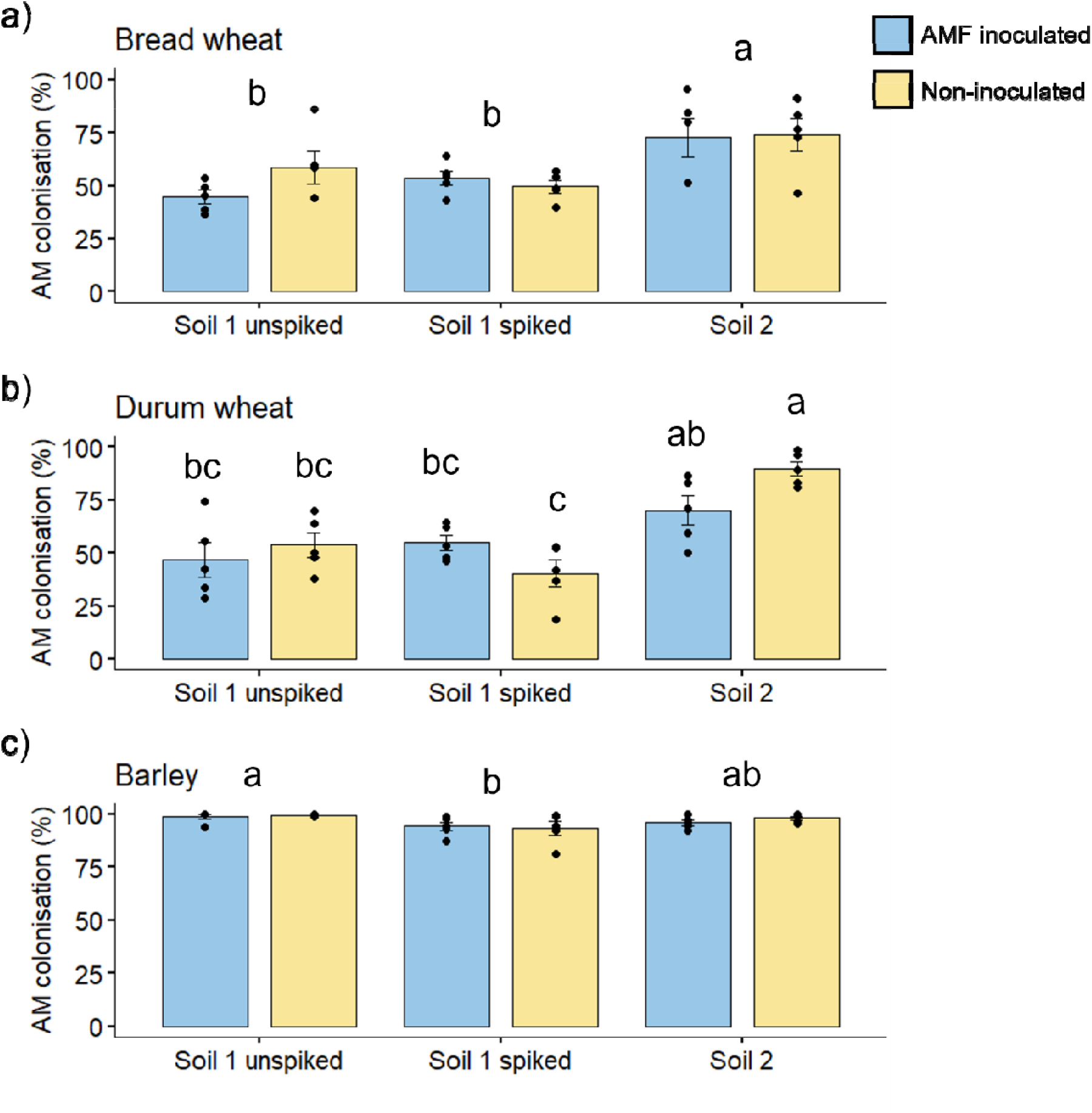
Percentage of root length colonised by arbuscular mycorrhizal (AM) fungi in bread wheat (a), durum wheat (b) and barley (c), either grown in Soil 1 unspiked, or spiked with PFAS, or Soil 2, naturally contaminated with PFAS, and inoculated (blue) or not (yellow) with a commercial source of the AM fungus *Rhizophagus irregularis* (Experiment 1). Bars are treatment mean ± standard error of the mean, *n*=5. Letters indicate differences following Tukey’s HSD *post hoc* test, where means that share a letter are not significantly different. Figures with three sets of letters indicate a significant main effect of *Soil*, while figures with six sets of letters indicate a significant interaction between *Soil* and *Mycorrhiza* in the ANOVA (see Table S2 Supporting information for further detail of statistical outcomes).

#### Plant biomass

In the bread wheat and barley plants, shoot biomass was affected by the Soil treatment only (Figure 2), where the plants grown in Soil 2 produced more shoot biomass than the respective crop when grown in Soil 1. In bread wheat, the plants in Soil 2 experienced a 25.3% increase in biomass compared to Soil 1 (mean across spiked and unspiked treatments), and in barley, there was a 17.2% increase in biomass in Soil 2. The durum wheat plants also displayed a small increase in biomass in Soil 2, but this was not statistically significant. There were no significant differences in biomass between the unspiked and spiked Soil 1.

**Figure 2.**
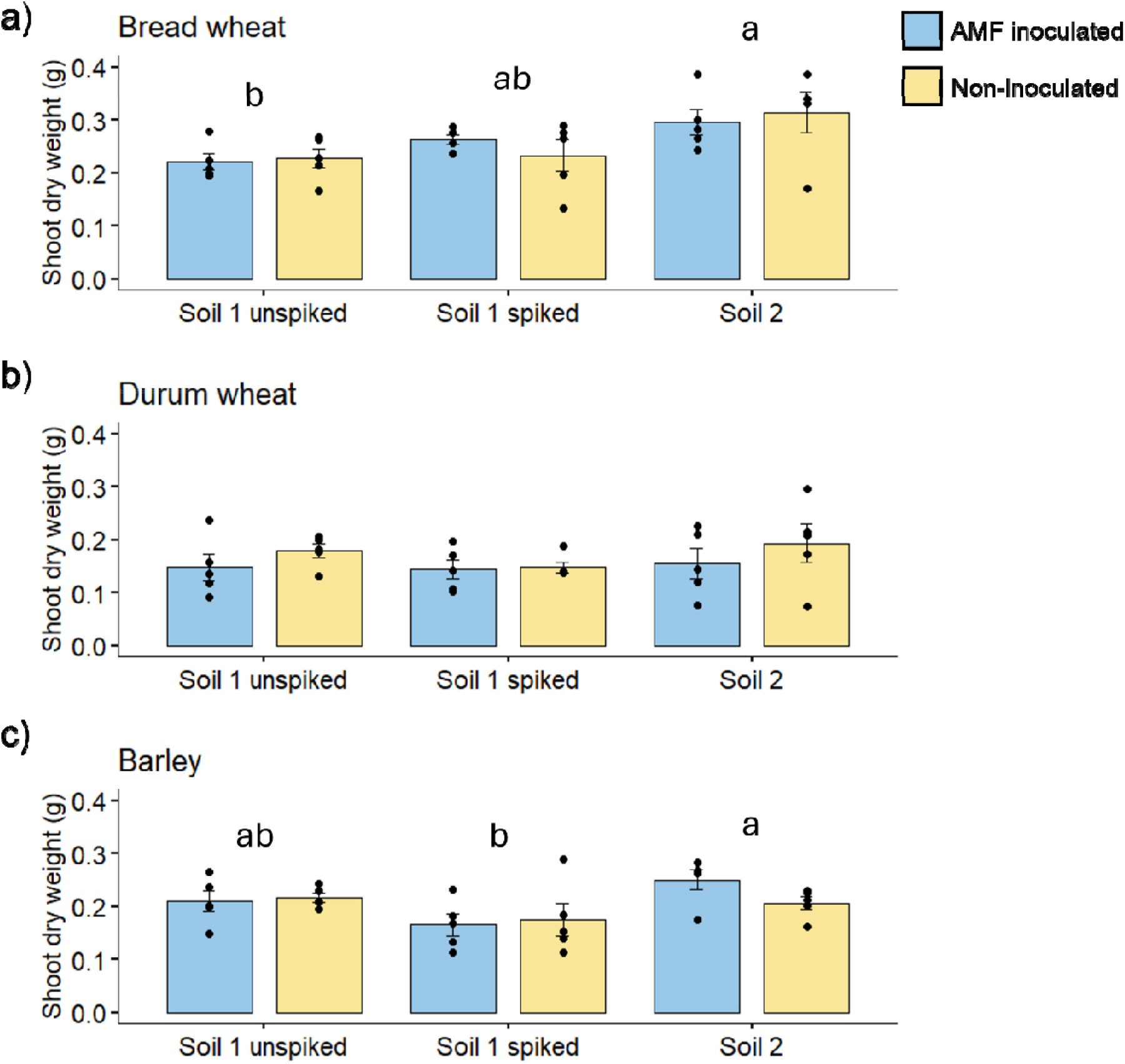
Shoot dry weights (g) of bread wheat (a), durum wheat (b) and barley (c), either grown in Soil 1 unspiked, or spiked with PFAS, or Soil 2, naturally contaminated with PFAS, and inoculated (blue) or not (yellow) with a commercial source of the AM fungus *Rhizophagus irregularis* (Experiment 1). Bars are treatment mean ± standard error of the mean, *n*=5. Letters indicate differences following Tukey’s HSD *post hoc* test, where means that share a letter are not significantly different. Figures with three sets of letters indicate a significant main effect of *Soil* in the ANOVA (see Table S2 Supporting information for further detail of statistical outcomes).

#### Shoot PFAS concentrations

The Experiment 1 plants were analysed for four perfluorosulfonic acid compounds: PFBS, PFPeS, PFHxS and PFOS. The concentration and the speciation profile of sulfonic acids in the plants was highly influenced by the soil they were grown in, either spiked with PFAS solution (Soil 1) or a PFAS contaminated soil (Soil 2) collected from the field (Figure 3).

**Figure 3.**
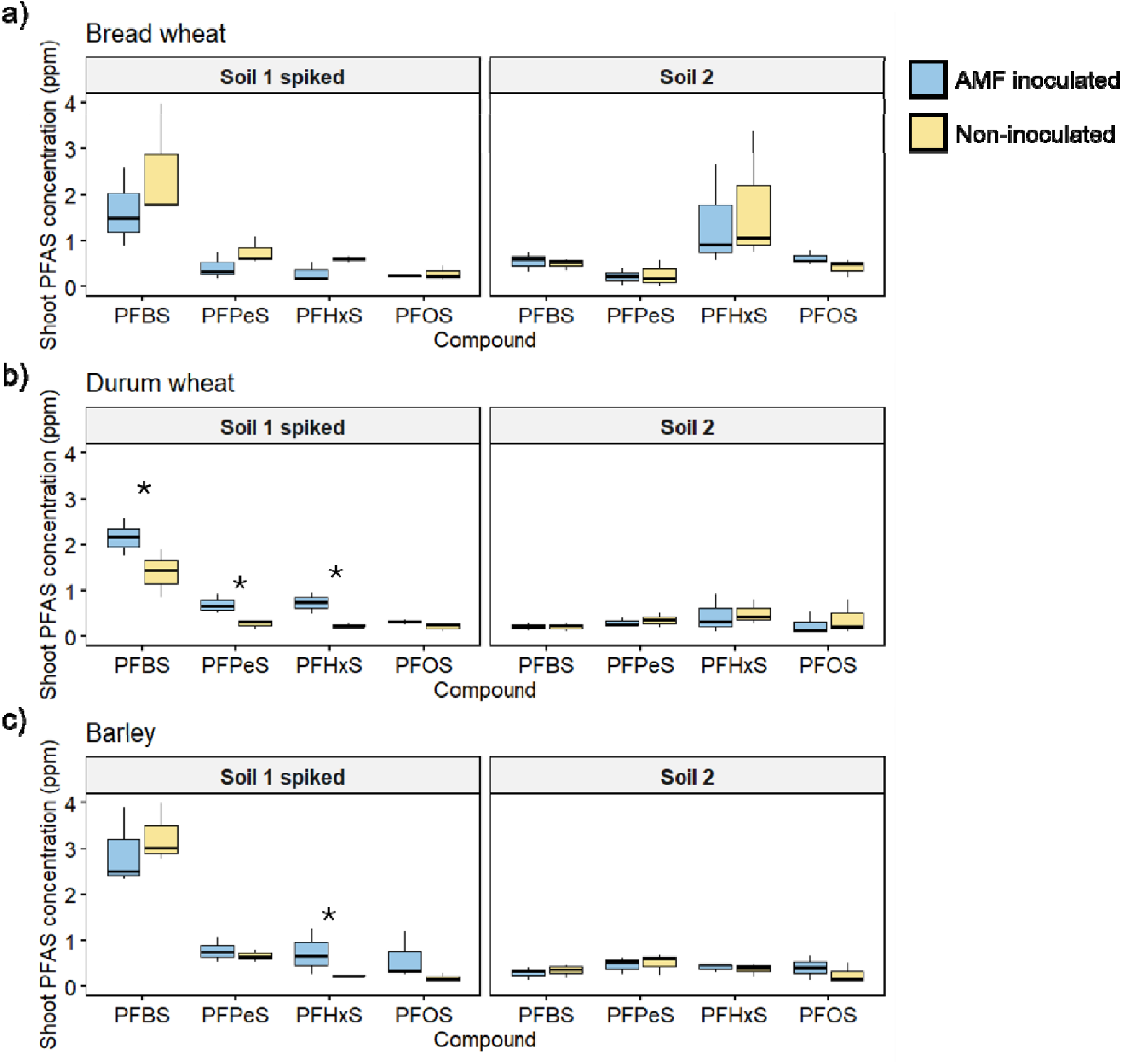
Concentration (ppm) of four different perfluorooctane sulfonic acids in the shoots of bread wheat (a), durum wheat (b) and barley (c), either grown in Soil 1 spiked with PFAS, or Soil 2, naturally contaminated with PFAS, and inoculated (blue) or not (yellow) with a commercial source of the AM fungus *Rhizophagus irregularis* (Experiment 1). Boxplots indicate the minimum, interquartile ranges, median (thick line), and maximum value, *n*=3. Asterisks indicate a significant difference in treatment mean between the AMF-inoculated and non-inoculated plants for an individual sulfonic acid, following Student’s *t*-test.

The durum wheat plants grown in the PFAS-spiked Soil 1 had greater shoot concentrations of PFBS, PFPeS and PFHxS when inoculated with the AM fungus *R. irregularis*. For example, shoot PFHxS concentration was mean 0.73 ppm in the inoculated durum wheat plants, compared to mean 0.25 ppm in the non-inoculated plants. In comparison, the durum wheat plants grown in Soil 1 that had not been spiked with PFAS had a mean PFHxS concentration of 0.01 ppm (Table S5 Supporting information). The barley plants had greater PFHxS concentration when inoculated with *R. irregularis*, but there were no significant effects of AM fungal inoculation on bread wheat.

In general, all three crop species accumulated higher concentrations of PFAS compounds in the spiked soil (Soil 1) than the naturally contaminated soil (Soil 2). In Soil 1, the PFAS compounds displayed the expected trend of higher concentrations of shorter chain compounds, but this was not seen in the plants grown in Soil 2, where PFBS concentration was low compared to the other measured compounds.

### Experiment 2

#### AM colonisation

In the second experiment, the plants were grown in Soil 1 that had been sterilised by autoclaving, and then was either re-inoculated with a commercial source of the AM fungus *R. irregularis*, or not inoculated. Thus, the extent of AM colonisation was greatly reduced in Experiment 2 compared to Experiment 1 which used Soil 1 that had not been sterilised. For example, in barley, the mean colonisation was 37.1% in Expt. 2, compared with 96.5% in the equivalent soil from Expt. 1 (Figure 4a). The bread wheat and durum wheat plants experienced relatively low AM colonisation (mean 9.1 % and 4.6% root length colonised, respectively) in the sterilised and re-inoculated Soil 1. In the sterilised, non-inoculated soil, none of the plants showed any evidence of AM colonisation under the microscope, confirming that the soil sterilisation process was effective.

**Figure 4.**
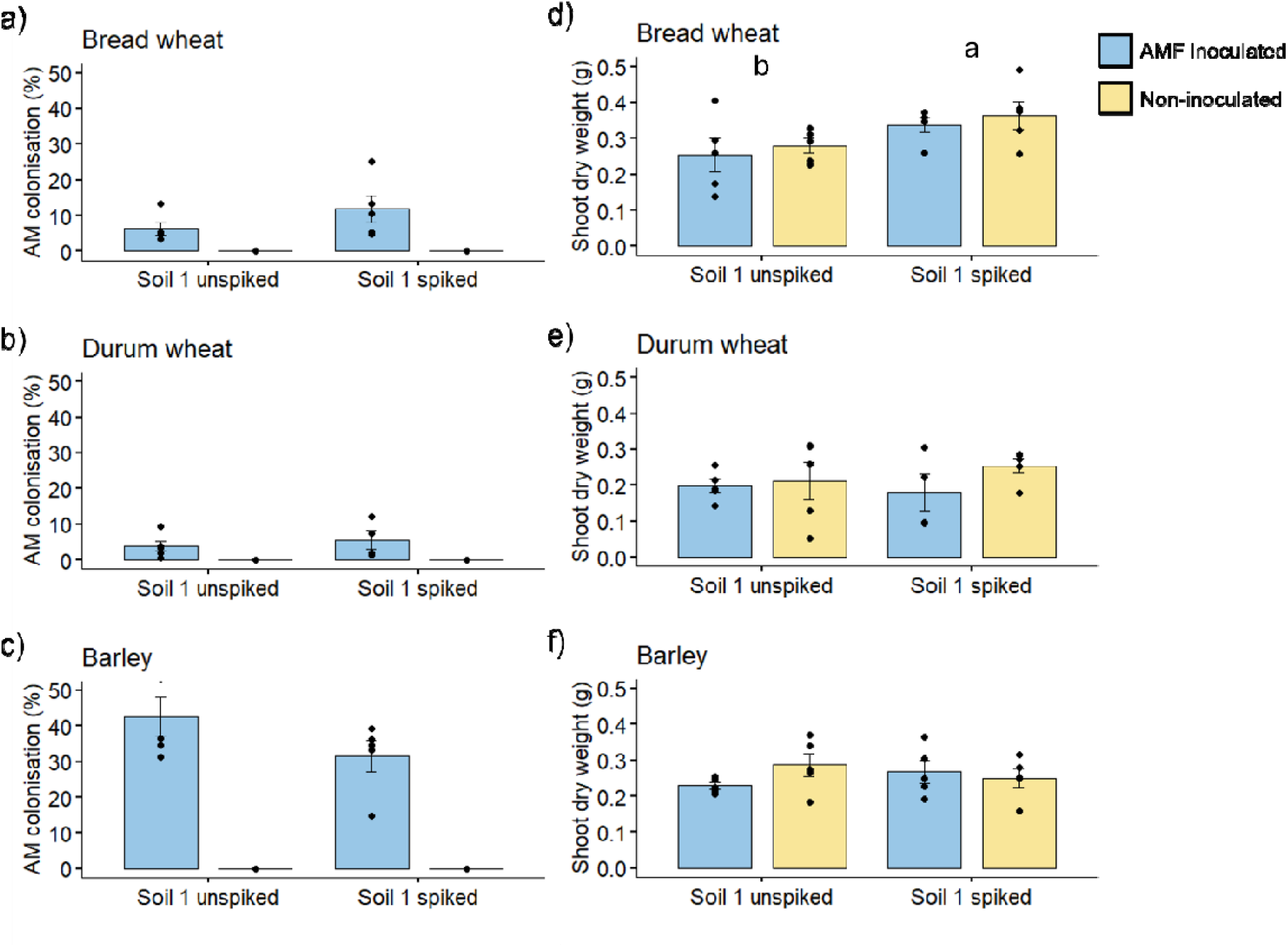
Percentage of root length colonised by arbuscular mycorrhizal (AM) fungi, and shoot dry weights of bread wheat (a,d), durum wheat (b,e) and barley (c,f), either grown in Soil 1 unspiked, or spiked with PFAS, and inoculated (blue) or not (yellow) with a commercial source of the AM fungus *Rhizophagus irregularis* (Experiment 2). Bars are treatment mean ± standard error of the mean, *n*=5. Letters indicate differences following Tukey’s HSD *post hoc* test, where means that share a letter are not significantly different. Figures with two letters indicate a significant main effect of *Soil* in the ANOVA.

#### Plant biomass

As with Expt. 1, the shoot biomass was not affected by AM colonisation in any of the three crop species (Figure 4b). For bread wheat, the shoot biomass increased significantly (an increase of 27%) when grown in the PFAS-spiked soil compared to the unspiked control soil. A similar but non-significant trend was observed in the bread wheat grown in Expt. 1 (increase of 10% in the spiked Soil 1). This “fertilisation effect” of the PFAS spiking of Soil 1 was not observed in the durum wheat or the barley.

#### Shoot PFAS concentrations

The plants from Expt. 2 were analysed for a subset of perfluourosulfonic (Figure 5) and perfluorocarboxylic (Figure 6) acid compounds. In general, the concentrations of the perfluorosulfonic acid compounds in durum wheat shoots in Expt. 2 were in the order of three times higher than in Expt. 1 (Figure 3), but the shoot dry weights were comparable between experiments. Furthermore, in this experiment, the durum wheat plants had significantly higher concentrations of every measured compound of PFAS when colonised by AM fungi. For example, shoot PFBS concentration was mean 4.12 ppm in the inoculated plants, compared to 1.65 ppm in the non-inoculated plants, and PFBA concentration was mean 3.14 ppm compared with 1.27 ppm.

**Figure 5.**
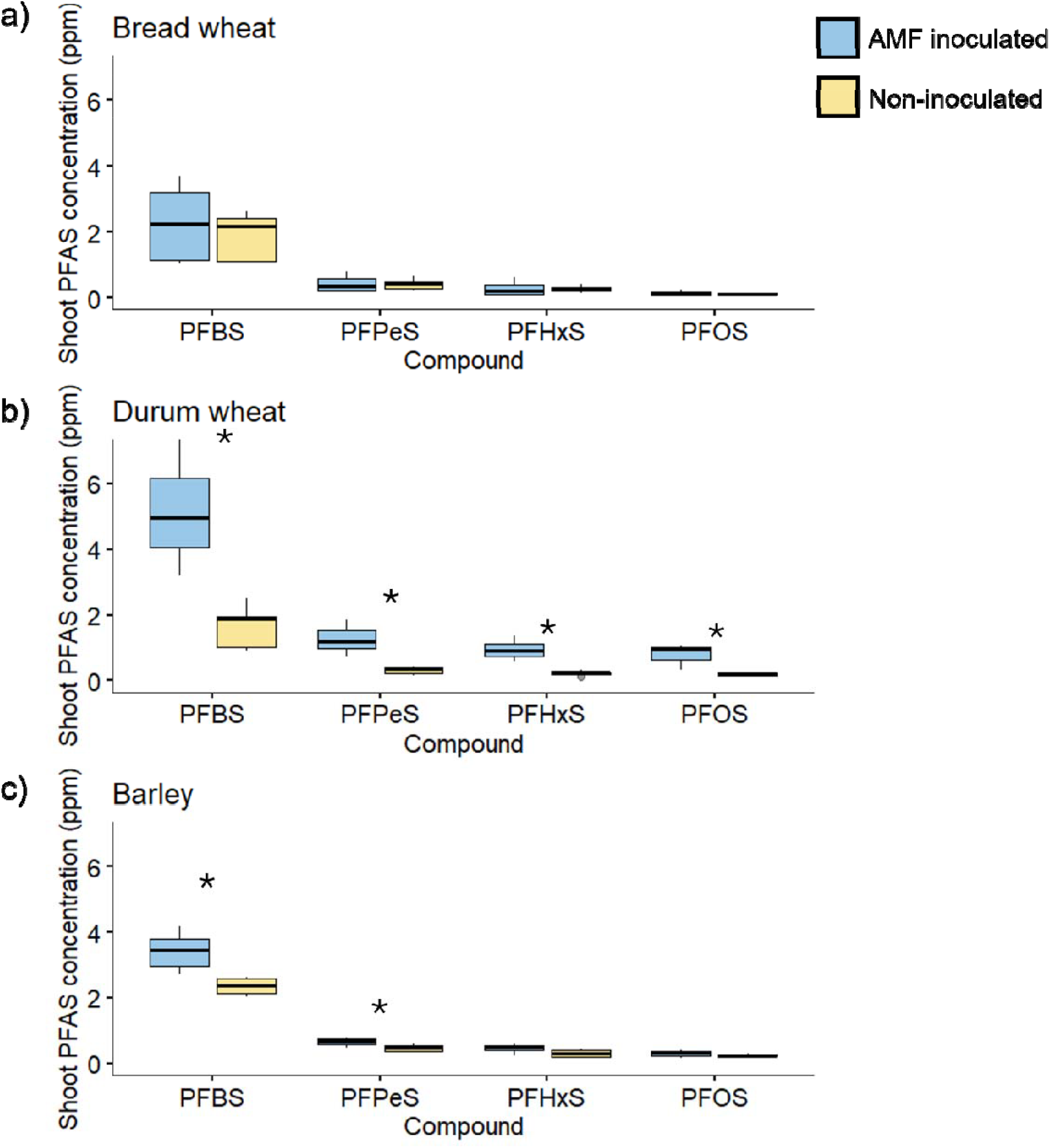
Concentration (ppm) of four different perfluorooctane sulfonic acids in the shoots of bread wheat (a), durum wheat (b) and barley (c), grown in Soil 1 spiked with PFAS, and inoculated (blue) or not (yellow) with a commercial source of the AM fungus *Rhizophagus irregularis* (Experiment 2). Boxplots indicate the minimum, interquartile ranges, median (thick line), and maximum value, *n*=5. Asterisks indicate a significant difference in treatment mean between the AMF-inoculated and non-inoculated plants for an individual sulfonic acid, following Student’s *t*-test.

**Figure 6.**
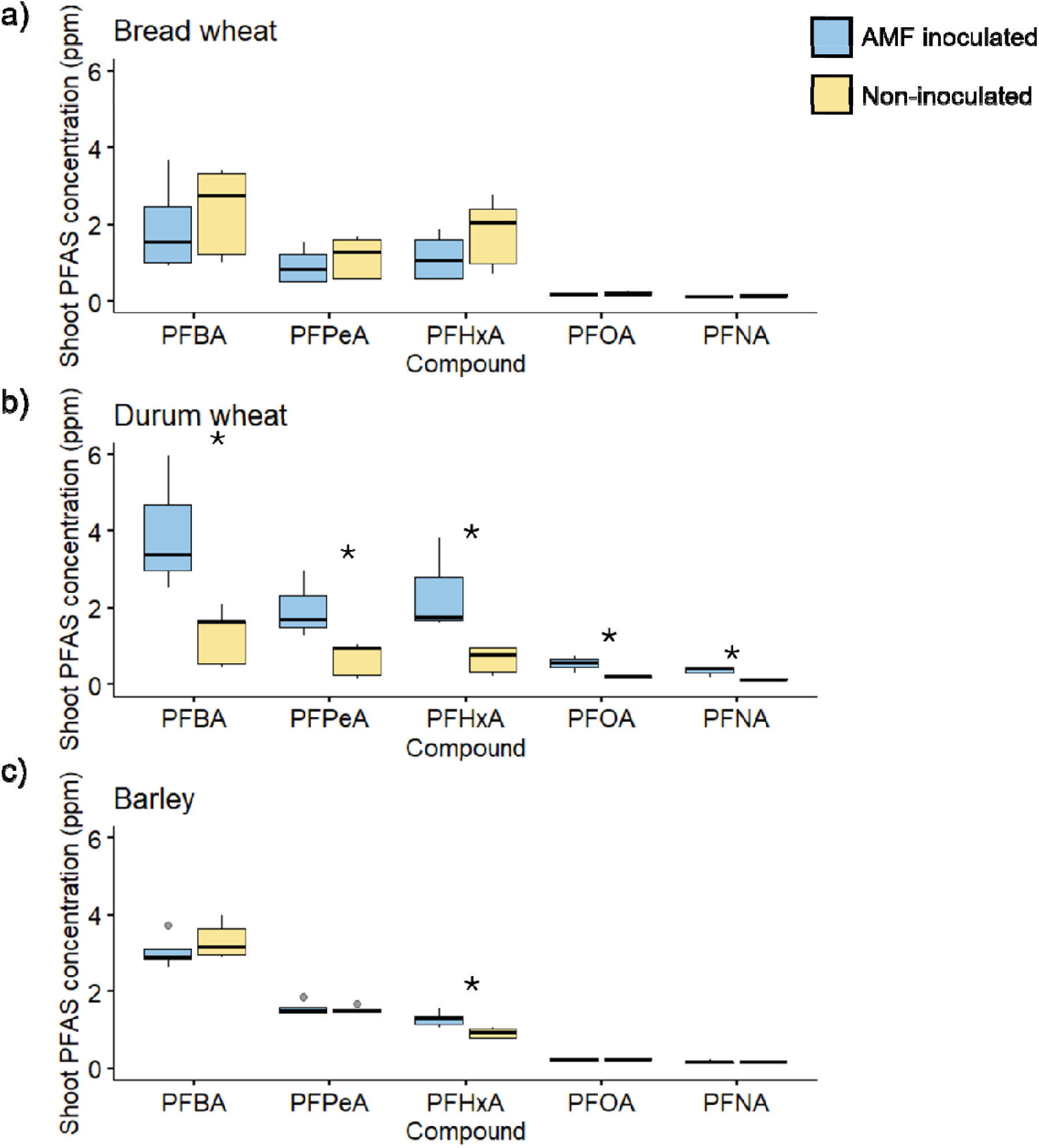
Concentration of five different perfluorocarboxylic acids in the shoots of bread wheat (a), durum wheat (b) and barley (c), grown in Soil 1 spiked with PFAS, and inoculated (blue) or not (yellow) with a commercial source of the AM fungus *Rhizophagus irregularis* (Experiment 2). Boxplots indicate the minimum, interquartile ranges, median (thick line), and maximum value, *n*=5. Asterisks indicate a significant difference in treatment mean between the AMF-inoculated and non-inoculated plants for an individual sulfonic acid, following Student’s *t*-test.

The barley plants also accumulated three measured PFAS compounds (two sulfonic, one carboxylic) more when colonised by AM fungi: PFBS, PFPeS and PFHxA. The PFBS concentrations in barley shoots were 2.26 ppm compared with 1.8 ppm.

In contrast to the other plant species, in bread wheat the concentration of PFAS compounds were generally lower than in durum wheat or barley, and there were no significant effects of AM colonisation on the concentration of any measured PFAS compounds.

The principal components analysis (PCA) was used to compare crops to each other in terms of their shoot PFAS concentrations. The associated biplot revealed that durum wheat was distinct in its PFAS concentration (generally indicated by principal component [PC] 1) and speciation (generally indicted by PC2) profile compared to barley and bread heat, which were not significantly different from each other (Figure 7). For example, durum wheat generally had higher concentrations of PFOS, PFOA and PFNA than the other two species.

**Figure 7.**
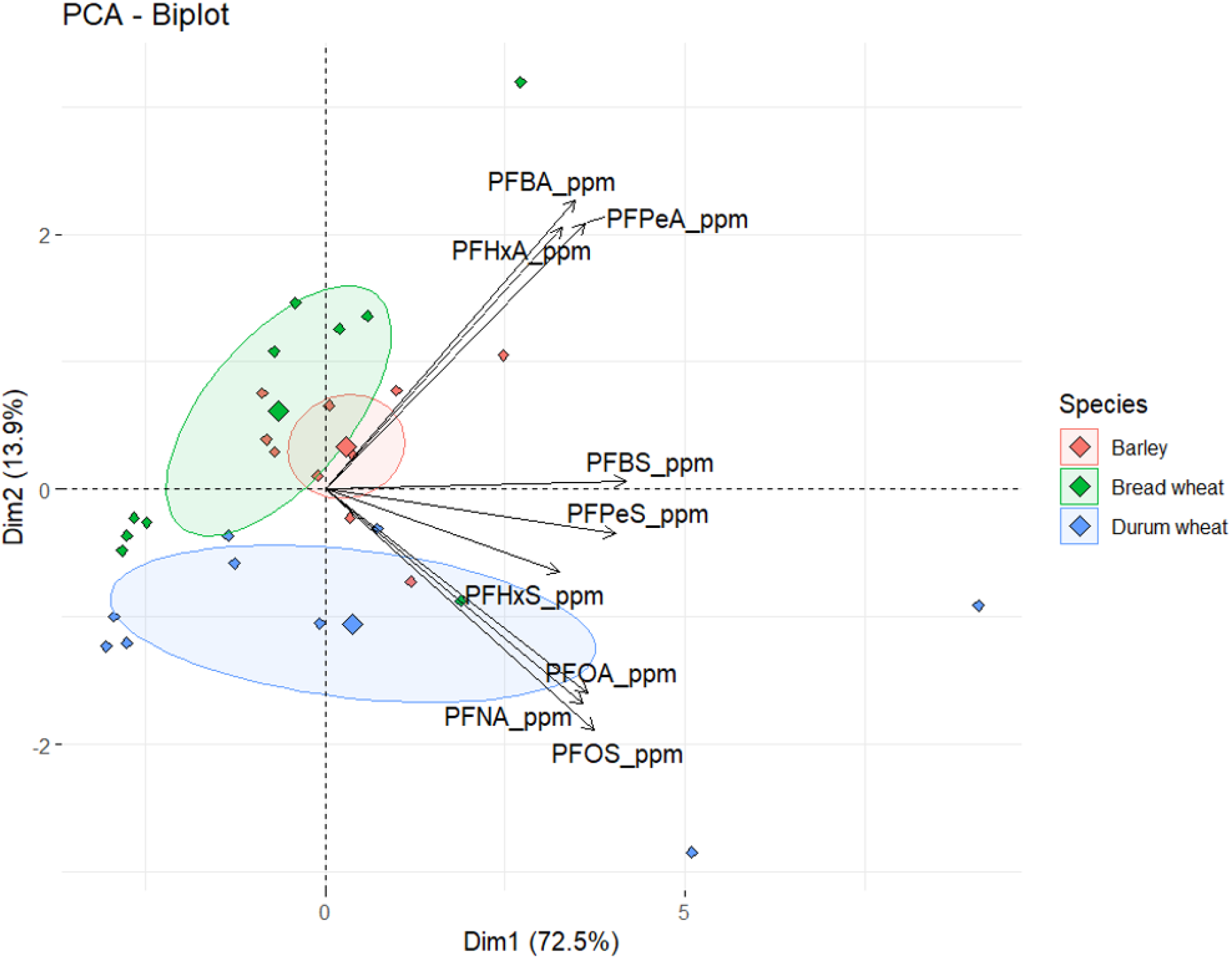
Principal components analysis (PCA) biplot displaying scores in the first two principal components (PC1: x-axis, PC2: y-axis) following PCA of the concentrations of nine PFAS compounds consisting of four perfluorooctane sulfonic acids and five perfluorocarboxylic acids in the shoots of barley (red), bread wheat (green) and durum wheat (blue) plants grown in a soil spiked with PFAS (Soil 1) (Experiment 2). The sign and magnitude of the contribution of variables is indicated by the loadings (arrows). The large diamonds signify the mean and 95 % confidence ellipse for each crop.

## Discussion

To our knowledge, this is the first study to empirically test the effect of AM fungi on the accumulation of various PFAS compounds in plant tissues. We tested the hypotheses that inoculation with AM fungi will have a: (1) neutral, (2) decreasing, or (3) increasing effect on shoot PFAS concentrations, and we tested these via two independent pot experiments that used non-sterilised soil (Expt. 1) or sterilised soil (Expt. 2); both experiments used the same soil, were spiked with PFAS in the same way, and included an inoculation treatment with a commercial source of *R. irregularis*.

### Cereal crops accumulate PFAS differently

The three cereal crops selected for these experiments (bread wheat, durum wheat and barley) are all important sources of food and/or feed throughout the world. Whilst they all belong to the Poaceae family, they exhibit diversity in many physiological traits, including their nutrition, protein, fibre and phenolics. It is therefore unsurprising that despite their genetic similarity, the three crops also exhibited differences in the magnitude and speciation of PFAS accumulation.

Here, the durum wheat plants generally had higher concentrations of longer chain PFAS compounds than barley or bread wheat, especially of PFOA (CF_2_,7), PFOS and PFNA (CF_2_,8). Barley on the other hand, tended to have higher concentrations of short-chain (CF_2_, 4-6) PFAS compounds. Longer-chain PFAS compounds, while less water soluble, have a greater tendency to bioaccumulate in biological tissues (Adu et al., 2023). Furthermore, they are presented at much higher concentration than other PFAS in our soils and most AFFF-contaminated soils. Others also observed that crop species can affect PFAS uptake by plants. In a field study conducted near a fluorochemical manufacturing facility, Liu et al. (2017 reported that the total concentration of 12 PFAS in wheat grains was more than 11 times higher than in maize grains.

While PFAS concentrations varied between the two soil types, the sandy nature of the soil may make the PFAS more readily available and prone to leaching. Previous work has shown that PFAS can leach almost completely, up to 100%, from sandy soils, regardless of their initial PFAS concentrations (Kabiri et al., 2022). However, these results were obtained under specific conditions involving leaching tests where soil was in contact with water at a liquid-to-solid (L/S) ratio of 1:20, subjected to vigorous shaking for 24 hours. Additionally, both our research and that of others have demonstrated that short-chain PFAS compounds leach more rapidly than long-chain PFAS in column leaching experiments. Due to their higher solubility and leachability, short-chain PFAS are also more bioavailable and more likely to be taken up by plants (Kabiri and McLaughlin, 2021; Kabiri et al., 2022; Stahl et al., 2013).

The elevated accumulation of longer chain compounds in durum wheat in this study may be due to its inherently higher protein content compared to bread wheat and barley (García-Puebla et al., 2023; Zilić et al., 2011). In Poland, a survey was undertaken of 89 commercial food products including flours, bread, pasta and noodles purchased from supermarkets and analysed for ten PFAS compounds (Surma et al., 2022); the authors found that among the processed foods (ie., not flours), the durum noodles (pasta) had a higher mean concentration of PFAS (5.6 ng g^-1^) than all other products, including wheat noodles (3.17 ng g^-1^). The tissue PFAS concentrations reached in our study were greater (by 2-3 orders of magnitude), but we measured PFAS in shoots rather than the food product where it would be diluted, and on much smaller plants than those the food products originated from.

More broadly, multiple studies have linked plant protein content with accumulation of PFAS compounds. In a plant study, it was found that root uptake and root-to-shoot translocation of PFOS and PFOA were positively correlated with plant protein content, so the legumes (soybean and mungbean) with higher protein contents, also had higher root concentration factors than the cereal crop ryegrass (Wen et al., 2016). More recently, Li et al. (2024 also found that soybean had the highest PFAS concentrations of the crops surveyed, including wheat and rice, reinforcing the correlation between protein content and PFAS accumulation in food crops.

The inclusion of a field contaminated soil in Expt. 1 served to highlight that PFAS accumulation in plants is not ubiquitous, and is highly dependent on the concentration and speciation of compounds in the soil, as well as soil chemical (Yang et al., 2024) and biological (Wu et al., 2023) properties that modify the solubility of the different compounds.

### AM fungal colonisation modulates shoot PFAS concentrations in a crop-dependent way

The relationship between AM fungi and their host plant hinges on the transfer of soil-based resources from the fungus to the host plant (Smith and Read, 2008). Generally, research into plant-AM fungal associations is focused on those resources that are necessary for plant growth and reproduction, such as a essential nutrients and water (Watts-Williams, 2022). However, anthropogenic soil contamination streams such as heavy metals and microplastics have more recently come in to focus as evidence builds that AM fungi can ‘protect’ the host plant from accumulating contaminants in their aboveground tissues (Chen et al., 2023; Gonzalez-Guerrero et al., 2008; Riaz et al., 2021). While it has been shown that AM fungi can physically prevent the uptake of microplastics into the tissues of host plants (Chen et al., 2023), there is limited published data on their relationship with plant PFAS uptake (Yan et al., 2025).

In this study, we found that inoculating durum wheat plants with *R. irregularis* led to an increase in the concentration of PFAS compounds in the shoots, in both sterile and non-sterile soils. This result was also observed to a lesser extent (ie., in fewer of the measured PFAS compounds) in the barley plants. The contrasting lack of any effect of AM fungi on the concentration of PFAS compounds in bread wheat suggests that the interaction between AM fungi and PFAS concentration effect is crop-dependent. There was no effect of AM colonisation on shoot biomass in these experiments, which is typical for cereal crops, and suggests it is unlikely that the observed effects on PFAS concentrations in durum and barley shoots were due to tissue dilution factors.

The effect of AM fungi on host plant PFAS accumulation may be direct (uptake by AM hyphae of PFAS from the soil and transport to plant across the peri-arbuscular membrane) or indirect (influence plant uptake via chemical modifications to the rhizosphere). On the other hand, the mechanisms of PFAS uptake by plants is incompletely understood. Some evidence suggests both passive and active transport pathways may be involved (Adu et al., 2023). Studies using metabolic and channel inhibitors indicate that PFOA and other PFAS can be taken up through energy-dependent processes, while PFOS uptake appears more passive, potentially involving aquaporins and anion channels. It has been proposed that wheat plants acquire PFAS from the soil by two mechanisms: ultra-short chain (CF_2_, 2 and 3) compounds that are highly water soluble could be taken up by aquaporins, while long-chain (CF_2_ ≥ 6) uptake is an energy-dependent active process (Zhang et al., 2019). Once there is more information available on the mechanism(s) for plant, and potentially fungal, PFAS uptake, it may help us predict which crop species may be more or less susceptible to tissue accumulation of PFAS.

### Soil PFAS contamination was not antagonistic to fungal or plant biomass

Soil contamination with heavy metals (e.g., cadmium, zinc), medical waste and radioactive sources have direct antagonistic effects on plant growth and reproduction. Meanwhile, in the cases of emerging contaminants such microplastics and PFAS, their accumulation in plant tissues may have little or no detrimental effect on the plant itself; instead, the concern lies in human and livestock consumption of contaminated plant material, particularly of long-chain PFAS compounds, which leads to bioaccumulation and, ultimately, detrimental health effects.

In our experiments, we did not observe any harmful effects of PFAS contamination on AM fungal colonisation of roots. In the case of barley, almost all roots were nearly 100% colonised by AM fungi including those exposed to PFAS in the spiked (with high concentration) and field-contaminated soils. This is in contrast to work that showed that AM fungal colonisation of spring onion roots was reduced in the presence of PFOS contamination in the soil (Yan et al., 2025). The difference was most likely due to the concentration of PFAS in the soil being around 10× higher than in our study. Additionally, Yan et al. (2025 measured arbuscular and vesicular colonisation, while we quantified the broader measure of total colonisation, which may have overlooked a PFAS-dependent effect on abundance of arbuscules or vesicles specifically.

We also observed no negative impacts on plant biomass of spiking a soil with PFAS, even though the shoots accumulated very high concentrations of some of the PFAS compounds. The capacity for herbaceous plants to accumulate PFAS in large amounts or to high concentrations in aboveground tissues is what makes them excellent candidates for PFAS phytoextraction, and warrants further research (Huff et al., 2020; Kavusi et al., 2023).

## Conclusions

We found that PFAS contamination in sandy soils had very limited effects on the extent of mycorrhizal colonisation in roots, or on plant biomass. However, the concentration of various PFAS compounds in the shoots was influenced by the AM status of the host plant, and the identity of the cereal crop. Durum wheat shoots consistently had higher concentrations of PFAS compounds when colonised by AM fungi, supporting the hypothesis that AM fungi promote PFAS uptake of the host plant, while there was no effect of AM fungi in bread wheat, which supports the null hypothesis. Barley was intermediate, with select PFAS compounds being elevated when the plant was mycorrhizal.

While this work has provided initial evidence that AM fungi can influence plant PFAS status under certain conditions, future studies of this nature should focus on using soils with PFAS concentrations and speciation that is realistic in an agricultural context, such as those with historical biosolid application.

## Acknowledgements

All authors acknowledge the Yitpi Foundation, South Australia for funding. SJWW and SK were each supported by Australian Research Council Discovery Early Career Researcher Awards (DE210100908, DE240100756). We thank Suhair Ahmed Hamad and Minshu Liang for excellent technical assistance. We thank Prof. Matthew Tucker for access to the barley seed, A/Prof. Stuart Roy for access to the bread wheat seed, and Australian Grains Technology for access to the durum wheat seed.

## References

Adu O, Ma X, Sharma VK. Bioavailability, phytotoxicity and plant uptake of per-and polyfluoroalkyl substances (PFAS): A review. Journal of Hazardous Materials 2023; 447: 130805.

Chen BD, Li XL, Tao HQ, Christie P, Wong MH. The role of arbuscular mycorrhiza in zinc uptake by red clover growing in a calcareous soil spiked with various quantities of zinc. Chemosphere 2003; 50: 839–846.

Chen H, Zhang X, Wang H, Xing S, Yin R, Fu W, et al. Arbuscular Mycorrhizal Fungi Can Inhibit the Allocation of Microplastics from Crop Roots to Aboveground Edible Parts. Journal of Agricultural and Food Chemistry 2023; 71: 18323–18332.

Cordner A, Goldenman G, Birnbaum LS, Brown P, Miller MF, Mueller R, et al. The True Cost of PFAS and the Benefits of Acting Now. Environ Sci Technol 2021; 55: 9630–9633.

Frew A, Powell JR, Heuck MK, Albornoz FE, Birnbaum C, Dearnaley JDW, et al. AusAMF: The Database of Arbuscular Mycorrhizal Fungal Communities in Australia. Global Ecology and Biogeography 2025; 34: e70090.

García-Puebla CA, Heredia-Olea E, López-Córdova JP, Dórame-Miranda RFs, Padilla-Torres CV, Rodríguez Félix F, et al. Use of durum wheat (Triticum Durum L.) with “yellow berry” as an alternative to malts in the production of ale-type beer: Physicochemical, quality of malts, and sensorial analysis. Journal of Cereal Science 2023; 109: 103613.

Giovannetti M, Mosse B. An evaluation of techniques for measuring vesicular arbuscular mycorrhizal infection in roots. New Phytologist 1980; 84: 489–500.

Gonzalez-Guerrero M, Melville LH, Ferrol N, Lott JNA, Azcon-Aguilar C, Peterson RL. Ultrastructural localization of heavy metals in the extraradical mycelium and spores of the arbuscular mycorrhizal fungus Glomus intraradices. Canadian Journal of Microbiology 2008; 54: 103–110.

Huff DK, Morris LA, Sutter L, Costanza J, Pennell KD. Accumulation of six PFAS compounds by woody and herbaceous plants: potential for phytoextraction. International Journal of Phytoremediation 2020; 22: 1538–1550.

Jha G, Kankarla V, McLennon E, Pal S, Sihi D, Dari B, et al. Per- and Polyfluoroalkyl Substances (PFAS) in Integrated Crop&Livestock Systems: Environmental Exposure and Human Health Risks. International Journal of Environmental Research and Public Health 2021; 18: 12550.

Kabiri S, McLaughlin MJ. Durability of sorption of per- and polyfluorinated alkyl substances in soils immobilized using common adsorbents: 2. Effects of repeated leaching, temperature extremes, ionic strength and competing ions. Science of The Total Environment 2021; 766: 144718.

Kabiri S, Tucker W, Navarro DA, Bräunig J, Thompson K, Knight ER, et al. Comparing the Leaching Behavior of Per- and Polyfluoroalkyl Substances from Contaminated Soils Using Static and Column Leaching Tests. Environmental Science & Technology 2022; 56: 368–378.

Kavusi E, Shahi Khalaf Ansar B, Ebrahimi S, Sharma R, Ghoreishi SS, Nobaharan K, et al. Critical review on phytoremediation of polyfluoroalkyl substances from environmental matrices: Need for global concern. Environmental Research 2023; 217: 114844.

Li X, Zhang B, Hou M, Qian C, Ji Z, Shi Y, et al. Occurrence of per- and polyfluoroalkyl substances in wheat, maize, rice, and soybean from chinese major grain producing regions. Journal of Hazardous Materials 2024; 480: 136509.

Liu Z, Lu Y, Shi Y, Wang P, Jones K, Sweetman AJ, et al. Crop bioaccumulation and human exposure of perfluoroalkyl acids through multi-media transport from a mega fluorochemical industrial park, China. Environment International 2017; 106: 37–47.

Nassazzi W, Lai FY, Ahrens L. A novel method for extraction, clean-up and analysis of per- and polyfluoroalkyl substances (PFAS) in different plant matrices using LC-MS/MS. Journal of Chromatography B 2022; 1212: 123514.

Ofoegbu PC, Wagner DC, Abolade O, Clubb P, Dobbs Z, Sayers I, et al. Impacts of perfluorooctanesulfonic acid on plant biometrics and grain metabolomics of wheat (Triticum aestivum L.). Journal of Hazardous Materials Advances 2022; 7: 100131.

Riaz M, Kamran M, Fang Y, Wang Q, Cao H, Yang G, et al. Arbuscular mycorrhizal fungi-induced mitigation of heavy metal phytotoxicity in metal contaminated soils: A critical review. Journal of Hazardous Materials 2021; 402: 123919.

Smith SE, Read DJ. Mycorrhizal symbiosis: Academic Press, New York, 2008.

Stahl T, Riebe RA, Falk S, Failing K, Brunn H. Long-Term Lysimeter Experiment To Investigate the Leaching of Perfluoroalkyl Substances (PFASs) and the Carry-over from Soil to Plants: Results of a Pilot Study. Journal of Agricultural and Food Chemistry 2013; 61: 1784–1793.

Surma M, Sznajder-Katarzyńska K, Wiczkowski W, Zieliński H. Assessment of Bioactive Surfactant Levels in Selected Cereal Products. Applied Sciences. 12, 2022.

Vierheilig H, Coughlan AP, Wyss U, Piche Y. Ink and vinegar, a simple staining technique for arbuscular-mycorrhizal fungi. Applied and Environmental Microbiology 1998; 64: 5004–5007.

Wang W, Rhodes G, Ge J, Yu X, Li H. Uptake and accumulation of per- and polyfluoroalkyl substances in plants. Chemosphere 2020; 261: 127584.

Watts-Williams SJ. Track and trace: how soil labelling techniques have revealed the secrets of resource transport in the arbuscular mycorrhizal symbiosis. Mycorrhiza 2022; 32: 257–267.

Watts-Williams SJ, Patti AF, Cavagnaro TR. Arbuscular mycorrhizas are beneficial under both deficient and toxic soil zinc conditions. Plant and Soil 2013; 371: 299–312.

Wen B, Li L, Zhang H, Ma Y, Shan X-Q, Zhang S. Field study on the uptake and translocation of perfluoroalkyl acids (PFAAs) by wheat (Triticum aestivum L.) grown in biosolids-amended soils. Environmental Pollution 2014; 184: 547–554.

Wen B, Wu Y, Zhang H, Liu Y, Hu X, Huang H, et al. The roles of protein and lipid in the accumulation and distribution of perfluorooctane sulfonate (PFOS) and perfluorooctanoate (PFOA) in plants grown in biosolids-amended soils. Environmental Pollution 2016; 216: 682–688.

Wu E, Wang K, Liu Z, Wang J, Yan H, Zhu X, et al. Metabolic and Microbial Profiling of Soil Microbial Community under Per- and Polyfluoroalkyl Substance (PFAS) Stress. Environmental Science & Technology 2023; 57: 21855–21865.

Yan H, Zhao T, Xu B, Lehmann A, Rillig MC. Arbuscular mycorrhizal fungi suffer from perfluorooctanoic acid in soil but ameliorate negative effects on plant performance. Applied Soil Ecology 2025; 212: 106169.

Yang H, Zhao Y, Chai L, Ma F, Yu J, Xiao K-Q, et al. Bio-accumulation and health risk assessments of per- and polyfluoroalkyl substances in wheat grains. Environmental Pollution 2024; 356: 124351.

Zhang L, Sun H, Wang Q, Chen H, Yao Y, Zhao Z, et al. Uptake mechanisms of perfluoroalkyl acids with different carbon chain lengths (C2-C8) by wheat (Triticum acstivnm L.). Science of The Total Environment 2019; 654: 19–27.

Zilić S, Barać M, Pešić M, Dodig D, Ignjatović-Micić D. Characterization of proteins from grain of different bread and durum wheat genotypes. Int J Mol Sci 2011; 12: 5878– 94.

