## Supporting information for "Inoculation with arbuscular mycorrhizal fungi can increase the concentration of PFAS compounds in cereal crops"

**Table S1.** Concentrations of PFAS compounds in Soil 1 (spiked) and Soil 2, prior to the experiments. MDL refers to method detection limit.

| PFAS concentration in mg/kg | Soil 1 spiked | Soil 2 | MDL |
| --- | --- | --- | --- |
| *Perfluoroalkyl Sulfonic Acids* |  |  |  |
| Perfluorobutane sulfonic acid (PFBS) | 0.109 | 0.002 | 0.0002 |
| Perfluoropentane sulfonic acid (PFPeS) | 0.119 | 0.005 | 0.0002 |
| Perfluorohexane sulfonic acid (PFHxS) | 0.550 | 0.058 | 0.0002 |
| Perfluoroheptane sulfonic acid (PFHpS) | 0.112 | 0.009 | 0.0002 |
| Perfluorooctane sulfonic acid (PFOS) | 5.725 | 1.760 | 0.0002 |
| Perfluorodecane sulfonic acid (PFDS) | 0.025 | 0.020 | 0.0002 |
| *Perfluoroalkyl Carboxylic Acids* |  |  |  |
| Perfluorobutanoic acid (PFBA) | 0.023 | 0.003 | 0.001 |
| Perfluoropentanoic acid (PFPeA) | 0.017 | 0.005 | 0.0002 |
| Perfluorohexanoic acid (PFHxA) | 0.120 | 0.021 | 0.0002 |
| Perfluoroheptanoic acid (PFHpA) | 0.040 | 0.004 | 0.0002 |
| Perfluorooctanoic acid (PFOA) | 0.087 | 0.011 | 0.0002 |
| Perfluorononanoic acid (PFNA) | 0.0005 | 0.003 | 0.0002 |
| Perfluorodecanoic acid (PFDA) | <MDL | 0.002 | 0.0002 |
| Perfluoroundecanoic acid (PFUnDA) | <MDL | 0.001 | 0.0002 |
| Perfluorododecanoic acid (PFDoDA) | <MDL | 0.001 | 0.0002 |
| Perfluorotridecanoic acid (PFTrDA) | <MDL | 0.0005 | 0.0002 |
| Perfluorotetradecanoic acid (PFTeDA) | <MDL | <MDL | 0.0005 |

**Table S2.** Gradient method used for PFAS analysis on Agilent 6495 LC/TQ

| Time (min) | A% | B% | Flow rate (min) |
| --- | --- | --- | --- |
| 0.00 | 98.00 | 2.00 | 0.400 |
| 0.20 | 98.00 | 2.00 | 0.400 |
| 10.00 | 5.00 | 95.00 | 0.400 |
| 12.00 | 5.00 | 95.00 | 0.400 |
| 12.50 | 98.00 | 2.00 | 0.400 |
| 15.00 | 98.00 | 2.00 | 0.400 |

**Table S3.** Triple quadrupole instrument conditions

|  | **Source Parameters** |
| --- | --- |
| Gas temperature | 200° C |
| Gas flow | 14 L/min |
| Nebulizer | 35 psi |
| Sheath Gas Temperature | 250° C |
| Sheath Gas Flow | 11 L/min |
| Capillary Voltage (Neg) | 3000 V |
| Nozzle Voltage (Neg) | 1500 V |

**Table S4.** ANOVA outcomes for plant traits in Experiments 1 and 2.

| *Expt. 1* |  |  |  |  |
| --- | --- | --- | --- | --- |
| AM colonisation |  | *Soil* | *Mycorrhiza* | *Soil * Mycorrhiza* |
|  | Bread wheat | **0.00184** | 0.465 | 0.344 |
|  | Durum wheat | <0.0001 | 0.403 | **0.0284** |
|  | Barley | **0.0203** | 0.693 | 0.663 |
| Shoot dry weight |  | *Soil* | *Mycorrhiza* | *Soil * Mycorrhiza* |
|  | Bread wheat | **0.00832** | 0.928 | 0.587 |
|  | Durum wheat | 0.484 | 0.209 | 0.746 |
|  | Barley | **0.0198** | 0.58 | 0.323 |
| *Expt. 2* |  |  |  |  |
| AM colonisation |  | *Soil* | *Mycorrhiza* | *Soil * Mycorrhiza* |
|  | Bread wheat | 0.205 | **0.000414** | 0.205 |
|  | Durum wheat | 0.658 | **0.00244** | 0.499 |
|  | Barley | 0.128 | **<0.0001** | 0.128 |
| Shoot dry weight |  | *Soil* | *Mycorrhiza* | *Soil * Mycorrhiza* |
|  | Bread wheat | **0.0232** | 0.446 | 0.969 |
|  | Durum wheat | 0.663 | 0.272 | 0.44 |
|  | Barley | 0.959 | 0.444 | 0.173 |

**Table S5.** Mean concentrations (ppm) of four perfluorosulfonic acid compounds in the shoots of bread wheat, barley and durum wheat grown in the unspiked Soil 1 (Experiment 1).

|  | PFBS | PFPeS | PFHxS | PFOS |
| --- | --- | --- | --- | --- |
| Bread wheat | 0.527 | 0.069 | 0.094 | 0.008 |
| Durum wheat | 0.069 | 0.013 | 0.011 | 0.009 |
| Barley | 0.483 | 0.116 | 0.075 | 0.026 |
